# Vegetally localized Vrtn functions as a novel repressor to modulate *bmp2b* transcription during dorsoventral patterning in zebrafish

**DOI:** 10.1101/119776

**Authors:** Ming Shao, Min Wang, Yi-Wen Ge, Yan-Jun Zhang, De-Li Shi

**Affiliations:** School of Life Science, Shandong University, 27 Shanda Nan road, Jinan 250100, China; Sorbonne Universités, UPMC Univ Paris 06, CNRS UMR7622, IBPS-Developmental Biology Laboratory, 75005 Paris, France

**Keywords:** BMP signalling, Bmp2b, Vrtn, vegetal maternal determinants, dorsoventral patterning, zebrafish

## Abstract

The vegetal pole cytoplasm represents a critical source of maternal signals for patterning the primary dorsoventral axis of the early embryo. Vegetally localized dorsal determinants are essential for the formation of the Spemann organizer, which expresses both bone morphogenetic proteins and their antagonists. The extracellular regulation of BMP signalling activity has been well characterized, however, transcriptional regulation of *bmp* genes along the dorsoventral axis remains largely unknown. Here, we report a novel mode of maternal regulation of BMP signalling in the lateral and ventral regions by analyzing the unexpected dorsalizing effect following vegetal yolk ablation experiment. We identified Vrtn as a novel vegetally localized maternal factor displaying dorsalizing activity. It functions as a novel transcriptional repressor to regulate *bmp2b* expression in the marginal region. Mechanistically, Vrtn binds *bmp2b* upstream sequence and inhibits its transcription independently of maternal Wnt/ß-catenin signalling. By creating *vrtn* loss-of-function mutation and analyzing maternal-zygotic mutant embryos, we further showed that Vrtn is required for the formation of dorsoventral axis. Our work thus unveils a novel maternal mechanism regulating BMP gradient during dorsoventral specification.

## INTRODUCTION

The formation of dorsoventral (DV) axis is one of the earliest developmental processes that establish the animal body plan. The dorsal determinants (DDs) model suggests an intimate relationship between the animal-vegetal polarity and the DV axis, because dorsalizing signals are enriched in the vegetal pole of the oocyte. Triggered by fertilization and controlled by microtubules, DDs are asymmetrically transported, from vegetal pole to the perspective dorsal side in fish and amphibians (Abrams and Mullins, 2009; Cuykendall and Houston, 2009; Mei et al., 2009; Nojima et al., 2010; Langdon and Mullins, 2011; Shao et al., 2012; Tran et al., 2012; Ge et al., 2014; Slack, 2014; Carron and Shi, 2016). Several candidate DDs including Dishevelled, GBP, Wnt11 in *Xenopus* (Miller et al., 1999; Weaver et al., 2003; Tao et al., 2005), and Wnt8a in zebrafish (Lu et al., 2011), have been suggested to be required for dorsal axis formation. In zebrafish, DDs activate Wnt/ß-catenin signalling in the dorsal region of the blastula, causing the stabilization and nuclear localization of ß-catenin2, which is essential for zygotic expression of Spemann organizer genes (Bellipanni et al., 2006). One of the pivotal evidences supporting the DDs model is the vegetal pole ablation (VAb) experiment. Removing vegetal yolk cell before the first cleavage reduces or blocks the activation of maternal Wnt/ß-catenin signalling and results in ventralization. This suggests that DDs first localize at the vegetal pole of the oocyte, and rapidly evacuate this region after fertilization (Mizuno et al., 1998; Ober and Schulte-Merker, 1999). Consistent with this model, several vegetally localized factors, including Syntabulin (Nojima et al., 2010) and Grip2a (Ge et al., 2014), have been shown to be involved in the dorsal transportation of DDs.

Bone morphogenetic proteins (BMPs), mainly zygotically supplied in zebrafish, are key factors required for ventral fate specification. Several studies have suggested that *bmp* genes initiate their zygotic transcription under the control of maternal factors. Gdf6a can activate zygotic *bmp* genes expression through its receptor Alk6 and/or Alk8 (Goutel et al., 2000; Sidi et al., 2003). In dorsalized maternal-zygotic (MZ) *spg* (MZ*spg*) or maternal (M) *spg* (M*spg*) embryos, in which *pou5f3* gene is mutated, the transcription of *bmp* genes is inhibited (Reim and Brand, 2006). In addition, maternal factors MGA, Max, and Smad4 act in the yolk syncytial layer (YSL) to maintain *bmp2b* expression and initiate a positive feedback loop of BMP signalling within the embryo (Sun et al., 2014). Thus the function of these factors is a prerequisite for zygotic expression of *bmp* genes in the lateral and ventral regions. In the dorsal region, however, initial zygotic *bmp2b* transcription is suppressed by DDs-mediated pathway (Leung et al., 2003). Weak expression then appears in the shield, and helps to generate a correct BMP activity gradient along the DV axis (Xue et al., 2014). The presence of BMP signalling in the Spemann organizer has been proposed as a mechanism that ensures the self-regulation of the morphogenetic field (Dosch and Niehrs, 2000; Reversade et al., 2005; De Robertis, 2006; Inui et al., 2012). This suggests that dorsal transcription of *bmp* genes is tightly regulated to adjust the activity of the Spemann organizer and pattern the DV axis. However, how the early transcription of *bmp* genes in other regions of the embryo is regulated remains largely elusive.

The zygotic transcription of *bmp* genes in the lateral and ventral regions is activated by different maternal factors (Goutel et al., 2000; Sidi et al., 2003; Reim and Brand, 2006), there are also clues for the presence of repressive factors that function independently of DDs to modulate their expression. In particular, VAb after 2-cell stage can produce unexpected dorsalization (Mizuno et al., 1998). These embryos displayed a general decrease of *bmp2b* expression, which was not accompanied with an increase in Wnt/ß-catenin signalling. This is intriguing but how it is achieved remains unclear. In this study, we investigated the mechanism underlying this novel mode of maternal regulation of *bmp2b* expression. Our findings indicate that the dorsalization caused by VAb after 2-cell stage occurs in the absence of the Spemann organizer, and is specifically associated with a disrupted expression pattern of *bmp2b* and a reduced BMP activity. Mechanistically, we identified and characterized a novel vegetally localized transcriptional repressor, Vrtn, which binds *bmp2b* upstream sequence and prevents its transcription. This work uncovers a novel dorsalizing signal that functions independently of DDs-related Wnt/ß-catenin signalling to regulate DV patterning.

## RESULTS

### VAb after 2-cell stage causes dorsalization independently of Wnt/ß-catenin signalling

In order to understand how VAb after 2-cell stage causes dorsalization, the effects were re-evaluated during early development, from 1- to 256-cell stages. We found that VAb-mediated dorsalization was stage-dependent. VAb between 2- to 16-cell stages (VAb@2cell) produced a high proportion of dorsalized embryos (Fig. S1A,B). They displayed enlarged dorsal tissues at the expense of ventral fate (Fig. S1C-E’). Some embryos showed the highest degree of dorsalization (C5) (Kishimoto et al., 1997), with radial neural marker expression, expanded paraxial tissues and absence of lateral tissues such as the perspective pronephric duct, as revealed by the expression pattern of *pax2.1, egr2b* (*krox20*) and *myoD* (Fig. S1E,E’). By contrast, VAb before 2-cell stage (VAb@1cell) mainly caused ventralization. The embryos showed increased pronephric precursor cells, reduced or absent paraxial and axial tissues (Fig. S1B,F-G’).

The dorsalized phenotype could be caused by activation of dorsalizing signals, such as maternal Wnt/ß-catenin, or by inhibition of ventralizing signals, such as BMP and zygotic Wnt/ß-catenin. An inhibition of zygotic Wnt/ß-catenin never produces a characteristic long and elliptical shape at 12 hpf, a phenotype reminiscent of BMP deficient embryos (Ramel et al., 2005). To see if VAb@2cell-induced dorsalization was due to activation of maternal Wnt/ß-catenin signalling, we examined target genes of this pathway, *goosecoid* (*gsc*), *dharma* (*dha*) and *chordin* (*chd*), at sphere or 50% epiboly stage by in situ hybridization (ISH). Compared to intact embryos, the expression of these genes was either normal or reduced (Fig. S2A-I). Consistently, the nuclear localization of endogenous ß-catenin at 3.5 hpf (hours post-fertilization) was unchanged or reduced (Fig. S2J-L), excluding the possibility that VAb@2cell triggers activation of Wnt/ß-catenin signalling.

### Cell non-autonomous inhibition of BMP signalling in VAb@2cell embryos

To test if VAb@2cell causes BMP inhibition, we first examined P-Smad1/5/8 level. Compared to intact embryos, a large majority of VAb@2cell embryos showed weak or absent P-Smad1/5/8 immunostaining (Fig. 1A). We also detected by western blotting an average of 42% decrease in P-Smad1/5/8 level after VAb@2cell, compared to a 20.6% increase after VAb@1cell (Fig. 1B,C). We then examined BMP target genes, *eve1* and *gata2*. The expression of *eve1* was reduced in a majority of VAb@2cell embryos at 50% epiboly (Fig. 1D), suggesting BMP inhibition. When the expression pattern of *gata2* and *gsc* was simultaneously examined at 70% epiboly, we found that 60% (*n*=15) of VAb@2cell embryos showed reduced *gata2* expression, but with varied *gsc* expression patterns. Notably, gata2 expression was inhibited in some VAb@2cell embryos with absence of *gsc* expression (Fig. 1E). In all VAb@2cell embryos examined (13/13), expression of the pan-mesoderm marker *no tail* (*ntl*) was unaffected (Fig. 1F). These suggest that BMP inhibition after VAb@2cell is independent of the Spemann organizer, and that the reduced expression of *eve1* and *gata2* is not due to a lack of mesoderm formation. Thus, there may be a maternal factor inhibiting BMP activity in the lateral and ventral regions.

**Figure 1.**
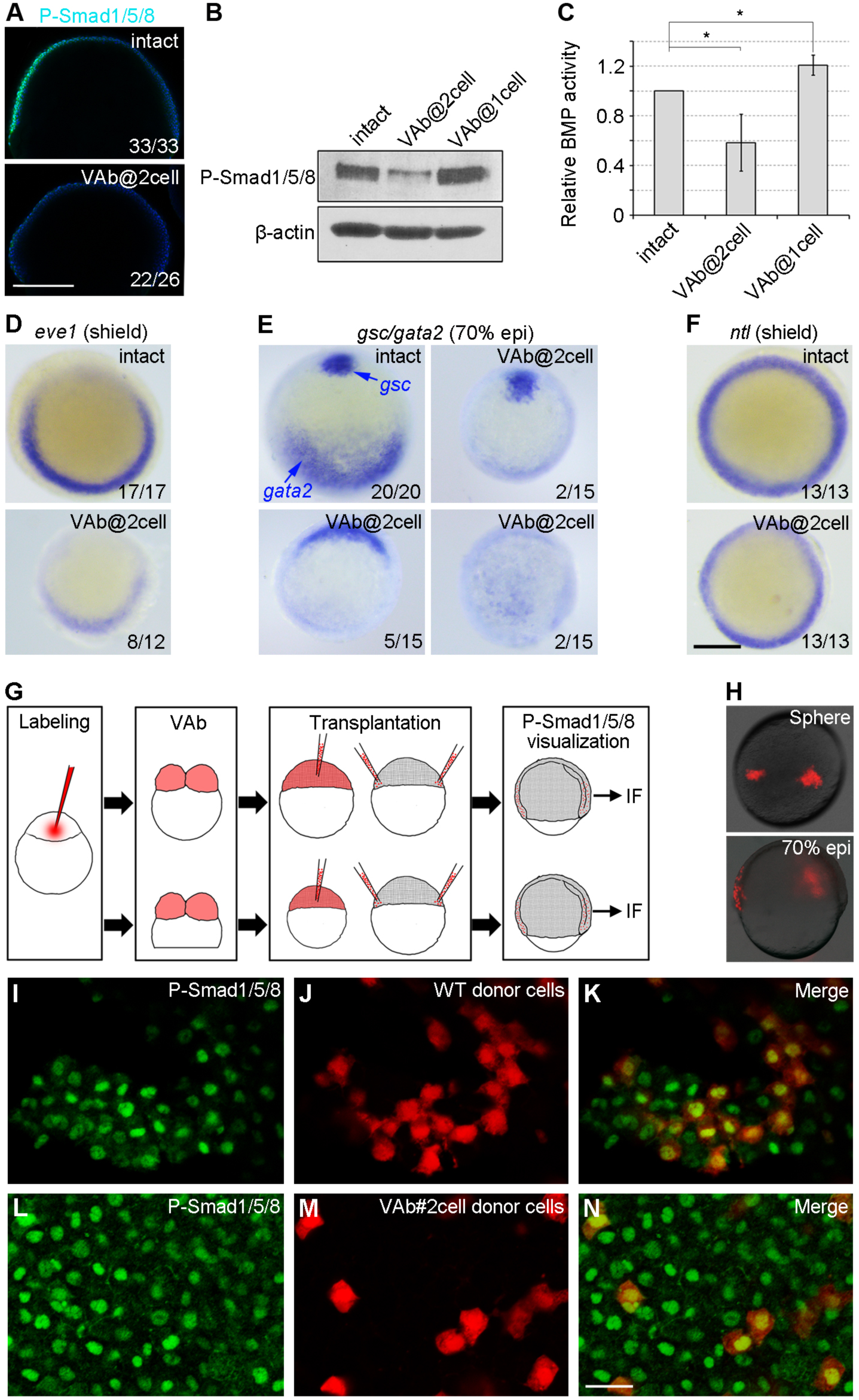
VAb@2cell inhibits BMP signalling. (A) Decreased P-Smad1/5/8 immunostaining (green), which is merged with DAPI (blue), in VAb@2cell embryos. (B,C) Western blotting and quantification of P-Smad1/5/8 level in VAb@2cell and VAb@1cell embryos at 70% epiboly. P-Smad1/5/8 level is normalized to ß-actin, and BMP activity in intact embryos is set as 1. Bars represent the mean values ± s.d. from three independent experiments. (D-F) In situ hybridization analysis of indicated genes. Animal pole view with dorsal region on the top. (G-N) Cell non-autonomous inhibition of BMP signalling in VAb@2cell embryos. (G) Procedure of cell transplantation experiment to visualize P-Smad1/5/8 by immunofluorescence (IF). (H) RLDx-labelled donor cells transplanted at different locations of the margin in unlabelled intact embryos at indicated stages. (I-N) Similar presence of P-Smad1/5/8 in transplanted cells derived from intact (I-K) or VAb@2cell (L-N) embryos. Scale bars: (A-F) 250 μm; (I-N) 20 μm.

We next performed transplantation experiments to examine how cells from VAb@2cell embryos interpret endogenous BMP signal. At sphere stage, deep blastoderm cells from RLDx-labelled VAb@2cell embryos were transplanted to two separate positions within the margin of unlabeled normal recipients to increase the probability of their ventral location (Fig. 1G,H). At 70% epiboly, P-Smad1/5/8 was similarly detected by immunostaining in transplanted cells from control and VAb@2cell embryos (Fig. 1I-N). Furthermore, when differentially labelled cells from intact and VAb@2cell embryos were mixed and transplanted to the ventral margin of unlabelled recipients at shield stage, they co-localized in posterior muscle, epidermis and blood (Fig. S3). These indicate that cells in VAb@2cell embryos were able to receive and activate BMP signal, and could normally undergo differentiation. Accordingly, injection of *bmp2b* mRNA (25 pg) in VAb@2cell embryos efficiently rescued the dorsalized phenotype (Fig. S4). Since BMP signalling in these embryos is inhibited in a cell non-autonomous manner, the putative BMP inhibitor should not be a component of the BMP transduction pathway.

### VAb@2cell inhibits zygotic bmp2b expression

One possibility is that the BMP inhibitor causes a deficiency of BMP ligands. In zebrafish, three BMP genes, *bmp2b, bmp7a* and *bmp4*, initiate their expression after mid-blastula transition, and maintain a gradient transcription during late blastula and gastrula stages. Both Bmp2b and Bmp7a are essential for DV patterning, but only Bmp2b/Bmp7a heterodimer is capable of activating BMP pathway (Little and Mullins, 2009). Consistently, loss of either *bmp2b* or *bmp7a* causes severe dorsalization (Dick et al., 2000; Dosch and Niehrs, 2000), while loss of *bmp4* only affects ventral fate in posterior tissues (Stickney et al., 2007). We thus examined by ISH the expression of these *bmp* genes in VAb@2cell embryos. Zygotic transcription of *bmp7a* and its expression in the early gastrula were unaffected (Fig. 2A-D). The expression of *bmp4* was also unchanged in the early gastrula (Fig. 2E,F), although it was not maintained at mid-gastrula stage (Fig. 2G,H), but this alone cannot account for the severely dorsalized phenotype. As for *bmp2b*, the initiation of zygotic transcription was unaffected (Fig. 2I,J), however, the marginal up-regulation from late blastula to early gastrula stages was strongly reduced (Fig. 2K,L,O). At mid-gastrula stage (75% epiboly), the expression of *bmp2b* in intact embryos was restricted to the ventral ectoderm in a relatively homogeneous pattern, whereas in more than half of VAb@2cell embryos, it became scattered in the ventral half (Fig. 2M-N’). Moreover, *bmp2b* expression was localized to blastoderm cells in intact embryos, in contrast to the internal YSL (I-YSL) in VAb@2cell embryos (Fig. 2M”,N”). By comparison, *bmp2b* expression was maintained or enhanced in VAb@1cell embryos (Fig. 2P-R). These indicate that VAb@2cell specifically inhibits *bmp2b* transcription in the margin.

**Figure 2.**
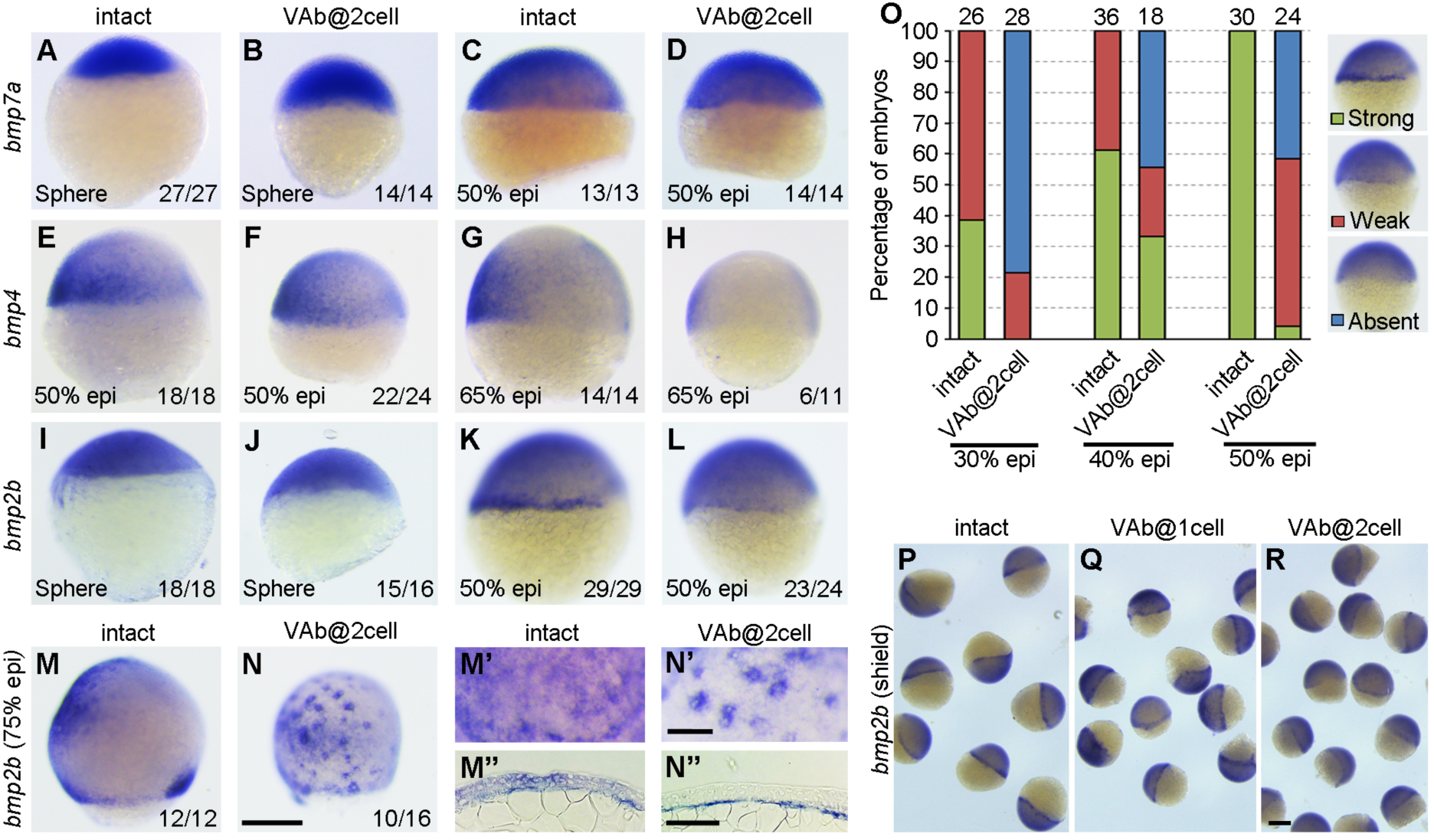
Differential expression of *bmp* genes in VAb@2cell embryos. (A-D) Similar expression pattern of *bmp7a* in intact and VAb@2cell embryos at indicated stages. (E-H) The expression pattern of *bmp4* remains unchanged in VAb@2cell embryos at 50% epiboly, but is strongly reduced at 65% epiboly. (I-L) The initiation of *bmp2b* expression is unaffected in VAb@2cell embryos at sphere-dome stage, but its zygotic transcription in the margin is reduced or absent at 50% epiboly. (M,N) Scattered *bmp2b* expression pattern in VAb@2cell embryos at 75% epiboly. Lateral views with dorsal on the right. (M’,N’) Higher magnification in the ventral-animal region of indicated embryos. (M’’,N’’) Localization of *bmp2b* in indicated embryos. (O) Statistical analysis of *bmp2b* marginal expression in intact and VAb@2cell embryos at indicated stages, with representative images shown on the right. Numbers on the top of each column indicate total embryos scored from two independent experiments. (P-R) Comparison of *bmp2b* expression in indicated embryos at 50% epiboly. The marginal *bmp2b* expression is maintained or enhanced in VAb@1cell embryos, but is suppressed in VAb@2cell embryos. Scale bars: (A-N, P-R) 250 μm; (M’,M”,N’,N”) 100 μm.

To further test whether blocking the marginal expression of *bmp2b* mimics the dorsalizing effect of VAb@2cell, we specifically inhibited *bmp2b* in the margin by injecting translation-blocking morpholino (*bmp2b*MO) in the yolk cell (2 ng/embryo) at 128-cell stage (Fig. S5A-C). The result showed that almost half of the injected embryos (*n*=45) were severely dorsalized (C4-C5). However, when *bmp2b*MO was injected in the YSL at 512-cell stage, nearly all embryos developed normally (Fig. S5B,E). Collectively, these results suggest that a maternal BMP inhibitory factor, which should repress *bmp2b* transcription in the margin, is responsible for the dorsalizing effect of VAb@2cell,

### Identification of Vrtn as a novel vegetally localized BMP inhibitor

Since VAb@2cell blocks BMP signalling, it is possible that a BMP inhibitor is vegetally localized like DDs and translocated away towards the animal pole during the first cell cycle. Although *dazl, celf1* and *mago nashi* (*magoh*) have been shown to be vegetally localized (Maegawa et al., 1999; Suzuki et al., 2000; Marlow and Mullins, 2008), their overexpression did not produce any dorsalizing effect (Fig. S6), suggesting that they could not function as BMP inhibitors. We thus sought to identify the factor(s) responsible for the dorsalizing and BMP-inhibiting activity in VAb@2cell embryos. Animal and vegetal pole regions from fertilized eggs were dissected and subjected to high throughput RNA-seq (Fig. 3A). In addition to a number of known factors, we uncovered *vrtn* as a novel vegetally enriched transcript (Table S1). This was further confirmed by ISH on fertilized eggs (Fig. 3B). As early as 2-cell stage, *vrtn* transcripts began to be unbiasly deposited as punctate forms at the margin of the blastodisc (Fig. 3C,D), suggesting a rapid transport. At 50% epiboly, *vrtn* transcripts were restricted to the entire margin and YSL (Fig. 3E,F). During gastrulation, they were maintained in the ventral margin but were down-regulated in the dorsal and animal pole regions (Fig. 3G). As development proceeds, *vrtn* expression shifted to the neural plate boundary and newly formed somites (Fig. 3,I), and was restricted in the tail-bud region (Fig. 3J). We then performed VAb at different stages to examine how this affects *vrtn* maternal expression. ISH at 32-cell stage showed that the marginal accumulation of *vrtn* transcripts was strongly reduced after VAb@1cell, but was essentially unaffected when VAb was performed after 2-cell stage (Fig. 3K). Analysis by qRT-PCR indicated that more than 60% of *vrtn* transcripts were removed after VAb@1cell, compared to a 20% reduction after VAb@2cell, and an absence of effect after VAb at 32-cell stage (Fig. 3L). Collectively, these observations reveal a vegetal localization and a rapid transport of maternal *vrtn* transcripts.

**Figure 3.**
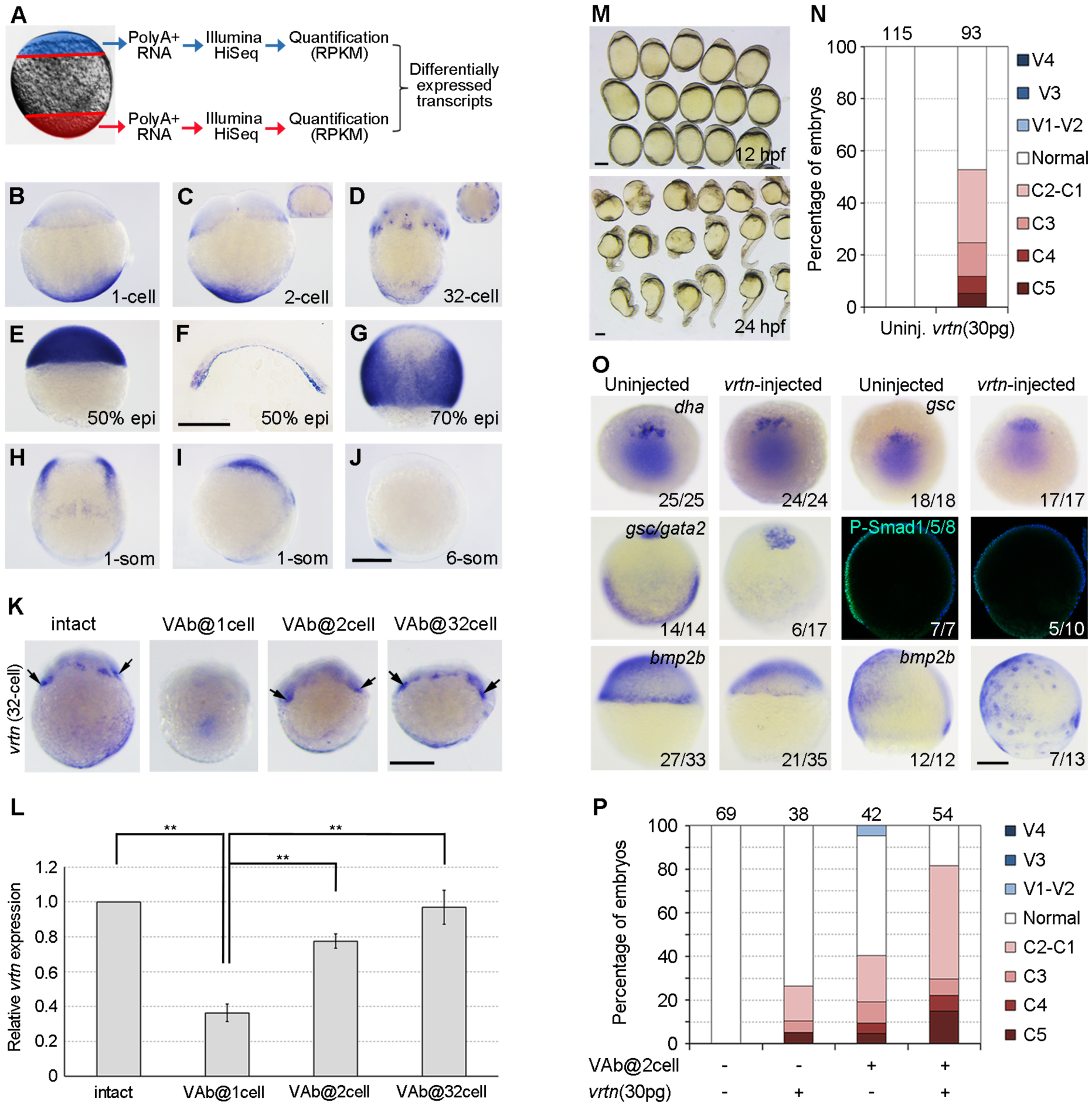
Vrtn overexpression suppresses BMP signalling. (A) Procedure for the identification of vegetal pole enriched transcripts. (B) Maternal *vrtn* transcripts are localized to the vegetal pole region in fertilized egg. (C) At 2-cell stage, *vrtn* transcripts are detected in the marginal region. The inset is an animal pole view. (D) At 32-cell stage, most *vrtn* transcripts are localized in the marginal region, as shown by the animal pole viewed inset. (E) Strong *vrtn* expression in the blastoderm at 50% epiboly. (F) Vertical section showing marginal and YSL expression of *vrtn* at 50% epiboly. (G) Dorsal view at 70% epiboly shows strong *vrtn* expression in the lateral region. (H,I) Dorsal (H) and lateral (I) views at 1-somite stage show *vrtn* expression in the neural plate boundary and forming somites. (J) Lateral view shows *vrtn* expression in the tail-bud at 6-somite stage. (K) In situ hybridization analysis of *vrtn* expression following VAb at indicated stages. Arrows denote the marginal accumulation of *vrtn* maternal transcripts. (L) qRT-PCR analysis of *vrtn* expression in indicated embryos. The relative expression level of *vrtn* in intact embryos is normalized as 1, and bars represent the mean values ± s.d. from three independent experiments. (M) Dorsalized embryos at 12 hpf and 24 hpf following Vrtn overexpression. (N) Statistical analysis of *vrtn*-dorsalized embryos at 24 hpf. Numbers on the top of each column indicate total embryos scored from three independent injection experiments. (O) Early effects of *vrtn* overexpression. The expression pattern of *dha* and *gsc* remains unchanged at sphere stage, and *gata2* expression at 70% epiboly and BMP activation at mid-gastrula stage are suppressed. The marginal transcription of *bmp2b* is inhibited at 50% epiboly, and scattered I-YSL expression of *bmp2b* is present at 70% epibloy. (P) Statistical analysis of the synergistic effect between VAb@2cell and *vrtn* overexpression. Numbers on the top of each column indicate total embryos scored from two independent experiments. Scale bars: 250 μm.

To test whether Vrtn functions as a BMP inhibitor, we performed overexpression by injecting the synthetic mRNA (100 pg) and found that this caused varied degrees of dorsalization, depending on the time of injection (Fig. 3M,N; Fig. S7). We then examined the expression of *dha* and *gsc*, and found that it was unaffected at sphere stage (Fig. 3O), indicating that the dorsalizing effect of Vrtn was independent of an ectopic activation of maternal Wnt/ß-catenin signalling. However, as in VAb@2cell embryos, *gata2* expression was inhibited in 35% of *vrtn*-injected embryos (*n*=17), while *gsc* expression was not obviously expanded (Fig. 3O). Consistently, P-Smad1/5/8 level was reduced in 50% (*n*=10), and *bmp2b* marginal expression was suppressed in 60% (*n*=35) of *vrtn*-injected embryos (Fig. 3O). In addition, 54% (*n*=13) of *vrtn*-injected embryos showed a scattered *bmp2b* expression pattern (Fig. 3O), as in VAb@2cell embryos. These strongly suggest that Vrtn overexpression mimics the effect of VAb@2cell. We further asked whether Vrtn could synergize with the dorsalizing effect of VAb@2cell. Embryos at 1-cell stage were injected with a low amount of *vrtn* mRNA (30 pg), and a fraction of these embryos was subjected to VAb@2cell. Intact embryos injected with *vrtn* and uninjected VAb@2cell embryos displayed 5.4% and 9.5% severely dorsalized phenotype (C4-C5), respectively. However, *vrtn*-injected VAb@2cell embryos exhibited 22.2% severely dorsalized phenotype, along with a significant increase of other less dorsalized phenotype (Fig. 3P). This clearly indicates a synergistic effect between Vrtn and VAb@2cell.

### VAb@2cell-caused dorsalization is suppressed in MZ*vrtn* mutant

We next used CRISPR/Cas9 to generate *vrtn* mutants. A 10 bp nucleotide deletion was obtained in the second exon. This produces a frame-shift in the 5’ region of the coding sequence and leads to a premature stop codon, which produces a truncated peptide with the first 17 amino acids in a total of 677 (Fig. 4A). In particular, it led to non-sense mediated decay. In MZ*vrtn* mutants, the *vrtn* transcripts were absent, especially in the vegetal pole region, although zygotic transcripts were detected at a much lower level (Fig. S8A-D). Analysis by qRT-PCR revealed a 22.7 folds decrease of the transcripts at cleavage stages and a 10.7 folds decrease at shield stage (Fig. S8E). Along with a rigorous allele-specific PCR analysis (Fig. S8F), it can be concluded that we have generated a loss-of-function mutation of *vrtn* gene.

**Figure 4.**
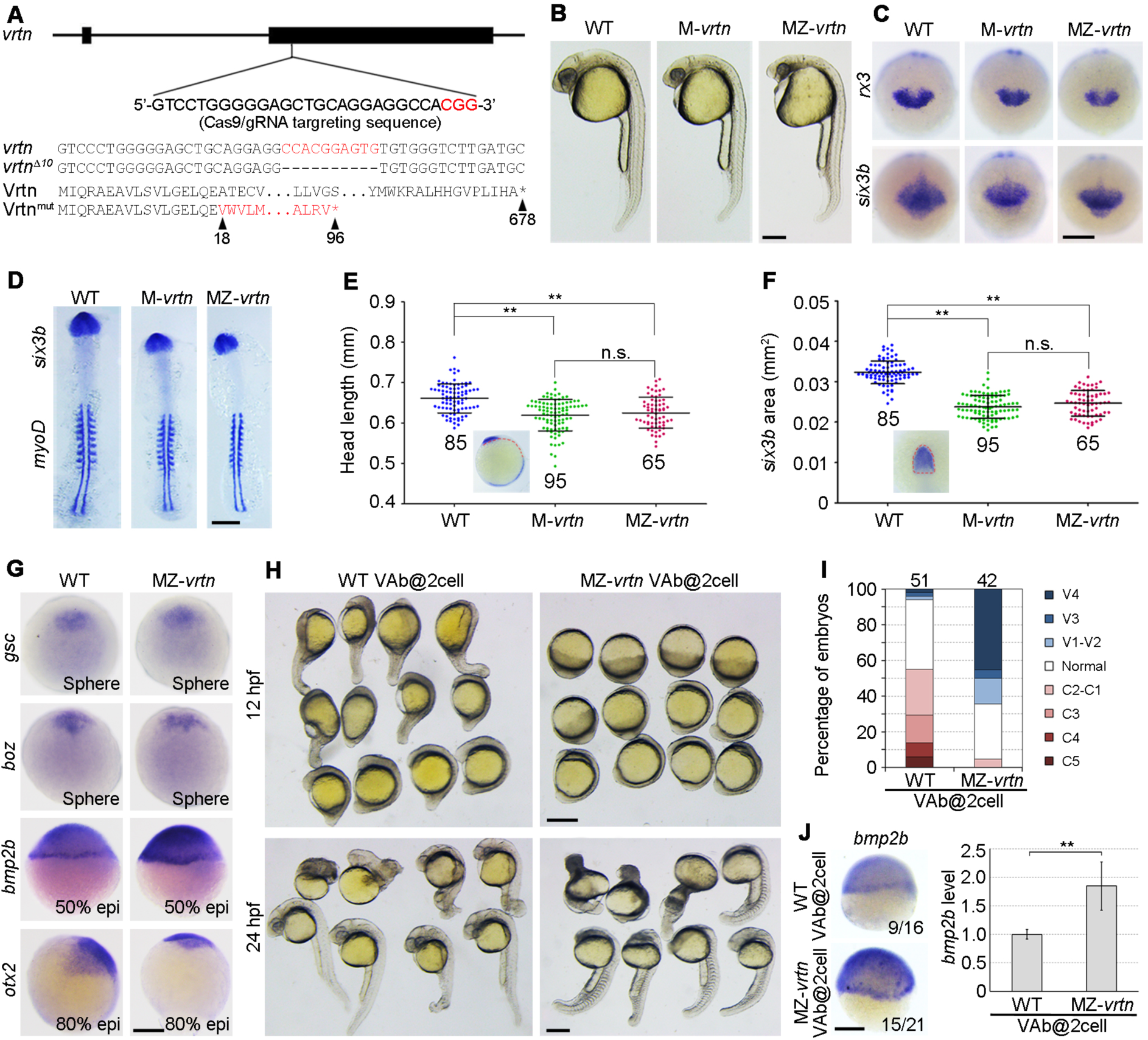
The dorsalizing effect of VAb@2cell is suppressed in MZ-*vrtn* mutant. (A) Targeted mutation of *vrtn* gene. The PAM region following the Cas9/gRNA targeting sequence is shown in red, and the mutated allele with the predicted protein sequence is indicated. (B) Representative images of indicated embryos at 27 hpf, showing smaller eyes and head size in *vrtn* mutants. (C) Reduced *rx3* and *six3b* expression domains in *vrtn* mutants at bud stage. (D) Expression pattern of *six3b* and *myoD* in flat-mounted embryos with equal somite number. (E,F) Analysis by scatter plots of head length and *six3b* expression area in indicated embryos at 6-somite stage. The red broken lines in the insets define the parameters of measurement. Horizontal lines represent the mean value ± s.d. from three batches of embryos with total numbers indicated. (G) The expression of *gsc* and *boz* remains unchanged at sphere stage in MZ-*vrtn* embryos (animal pole views), while the expression of *bmp2b* is enhanced at 50% epiboly, and the expression of *otx2* is reduced at 80% epiboly (lateral views with dorsal on the right). (H) *MZ-vrtn* embryos are ventralized following VAb@2cell. (I) Statistical analysis of the ventralizing effect. Numbers on the top of each column represent total embryos scored in three independent experiments. (J) In situ hybridization and qRT-PCR analyses of *bmp2b* expression at shield stage in WT and *MZ-vrtn* embryos subjected to VAb@2cell. The embryos are shown as lateral views. The average *bmp2b* expression level in WT VAb@2cell embryos is set as 1 using *ß-actin* as a loading control, and bars represent the mean values ± s.d. from six independent experiments. Scale bars: 250 μm.

The M*vrtn* and MZ*vrtn* embryos displayed a ventralized phenotype with a small head at 27 hpf (Fig. 4B), similarly as VAb@1cell embryos. However, the ventralization in VAb@1cell embryos is more severe probably due to the simultaneous removal of DDs. To exclude the possibility that the reduction in head size may be caused by a developmental delay of mutant embryos, we first visualized the eye field and the forebrain at bud stage by ISH using *rx3* and *six3b*, respectively, and found an evidently reduced expression domain of these genes in *Mvrtn* and MZ*vrtn* mutants (Fig. 4C). We further labelled somite stage embryos with *six3b* and *myoD*, and measured head length and *six3b* expression domain. The result showed that the head size was significantly reduced in *Mvrtn* and MZ*vrtn* mutants compared to wild-type (WT) embryos with an equal number of somites, while no difference was observed between *Mvrtn* and MZ*vrtn* embryos (Fig. 4E,F). These indicate that the lack of maternal *vrtn* is sufficient to generate ventralized phenotype. ISH analysis further indicated that *gsc* and *boz* expression was unaffected at mid-blastula stage, but *bmp2b* expression was enhanced at 50% epiboly, and *otx2* expression domain was reduced at 80% epiboly (Fig. 4G).

To analyse if the dorsalizing effect of VAb@2cell depends on Vrtn activity, we compared the phenotypes following VAb@2cell in WT and MZ*vrtn* embryos. In sharp contrast to VAb@2cell WT embryos, almost no dorsalization was observed in VAb@2cell MZ*vrtn* embryos (Fig. 4H). Intriguingly, they displayed essentially ventralized phenotypes (Fig. 4I), with strongly increased *bmp2b* expression at shield stage as revealed by ISH and qRT-PCR (Fig. 4J). This suggests that early *bmp2b* transcription is tightly regulated by Vrtn, and that MZ*vrtn* embryos are more sensitive to the reduction of DDs. To directly test this possibility, we injected *ß-catenin2* morpholino (*ß-cat2*MO) into WT and MZ*vrtn* embryos to compare the ventralizing effect. As reported (Bellipanni et al., 2006), most injected WT embryos exhibited mildly ventralized phenotypes.

However, in injected MZ*vrtn* embryos, there was a significant increase of severely ventralized phenotype (Fig. S9). This implies that DDs and Vrtn functionally interact and cooperate to control the BMP morphogenetic field.

The mildly ventralized MZ*vrtn* mutant phenotype with respect to the dorsalizing effect of VAb@2cell in WT embryos may be interpreted by the presence of other DDs-independent vegetal factors that are transported more slowly to the animal pole and function to counteract the activity of Vrtn. To test this possibility, we extracted total vegetal RNA at 2- to 4-cell stages in MZ*vrtn* embryos, which avoids the interference by endogenous Vrtn protein, and coinjected with *vrtn* mRNA (250 pg) in WT embryos. Using P-Smad1/5/8 as a readout, we found a 60% decrease of BMP activity in vrtn-injected embryos, and a significant rescue by the vegetal RNA, which did not affect BMP activity when injected alone (Fig. 5A,B). To examine how Vrtn activity may be temporally regulated by the putative vegetal factor(s), we assayed vegetal RNA from MZ*vrtn* embryos at later stages for their activity to antagonize Vrtn-mediated dorsalization. We reproducibly observed a robust rescue of P-Smad1/5/8 level only with 4-cell stage vegetal RNA (Fig. 5C,D). Accordingly, the Vrtn-dorsalized phenotype was also efficiently alleviated (Fig. 5E,F). These observations establish that Vrtn represents a novel vegetally localized factor that functions as a dorsalizing signal by inhibiting *bmp2b* transcription; they also suggest that Vrtn activity might be antagonized by other vegetal factors with a specific kinetics of animal pole transport.

**Figure 5.**
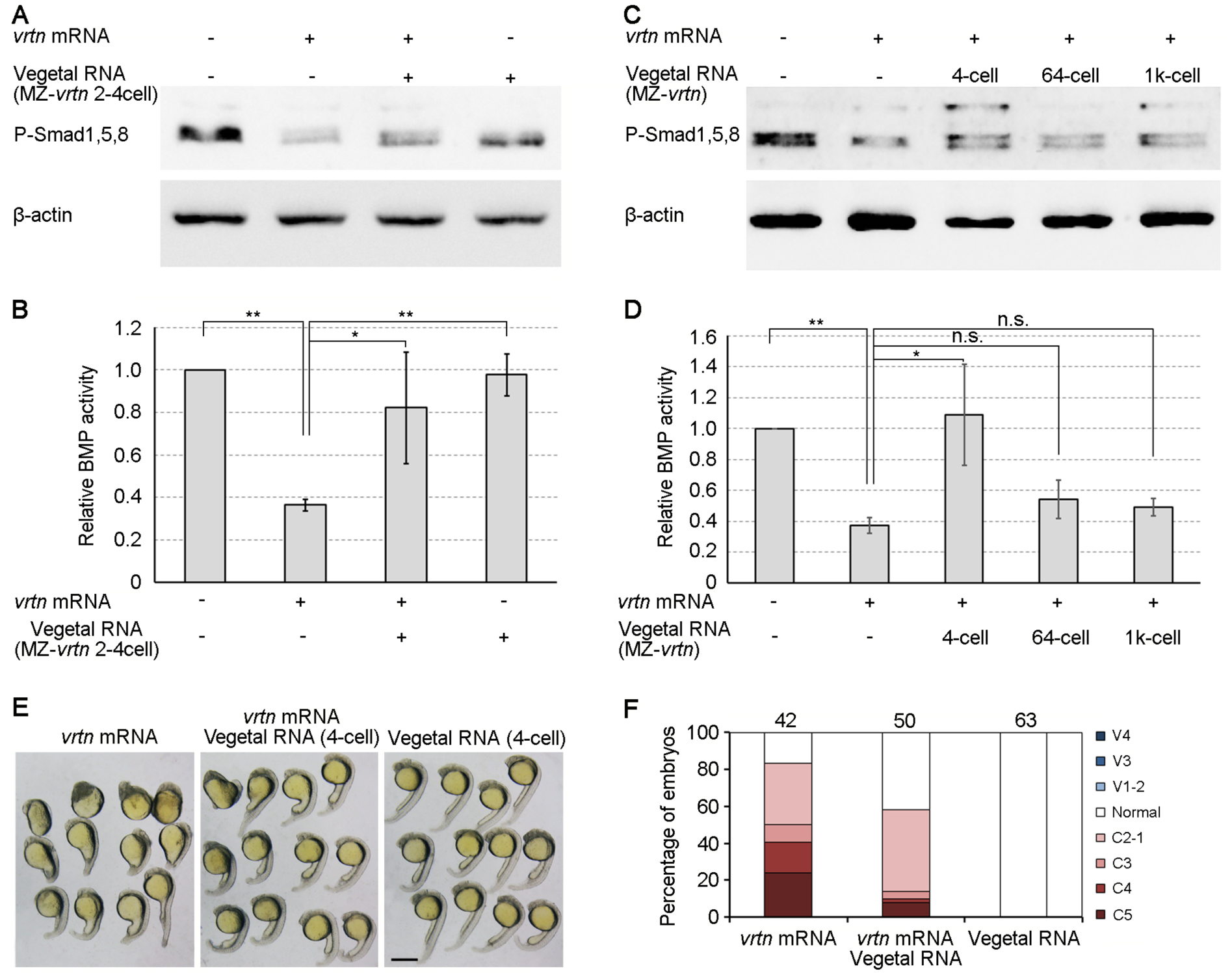
Vegetal RNA antagonizes the dorsalizing and BMP-inhibiting activity of Vrtn. (A) Western blotting at 50% epiboly shows that vegetal RNA from 2-cell to 4-cell stage rescues P-Smad1/5/8 level inhibited by Vrtn overexpression. (B) P-Smad1/5/8 level is normalized to ß-actin and quantified with respect to the BMP activity in uninjected embryos. Bars represent the mean value ± s.d. from four independent experiments. (C) Western blotting shows the temporal regulation of Vrtn activity by vegetal RNA. (D) Quantification of P-Smad1/5/8 level indicates that vegetal RNA from 64-cell or 1k-cell stage has no effect on Vrtn activity. Bars represent the mean value ± s.d. from three independent experiments. (E) Representative 24 hpf embryos showing the rescue of Vrtn-dorsalized phenotype by vegetal RNA. (F) Statistical analysis of the rescuing effect. Numbers on the top of each column indicate total embryos scored from two independent experiments. Scale bar: 200 μm.

### Vrtn acts as a transcriptional repressor

To further understand how Vrtn functions, its subcellular localization in zebrafish cells was analysed by immunostaining. Consistent with the nuclear localization in human cell lines (Mikawa et al., 2011), myc-tagged zebrafish Vrtn was enriched in the nucleus, but was released to the cytoplasm in M-phase cells (Fig. 6A,B). Since no functional domain could be predicted on the protein, we aligned 47 Vrtn sequences from 45 species and identified five conserved regions, designated as CD1 to CD5 (Fig. S10). These domains were individually deleted and overexpressed by injecting the corresponding mRNA (100 pg). The results showed that deletion of CD1 or CD3 severely disrupted Vrtn nuclear targeting (Fig. S11), suggesting that potential nuclear localization signals may be located separately in the protein.

**Figure 6.**
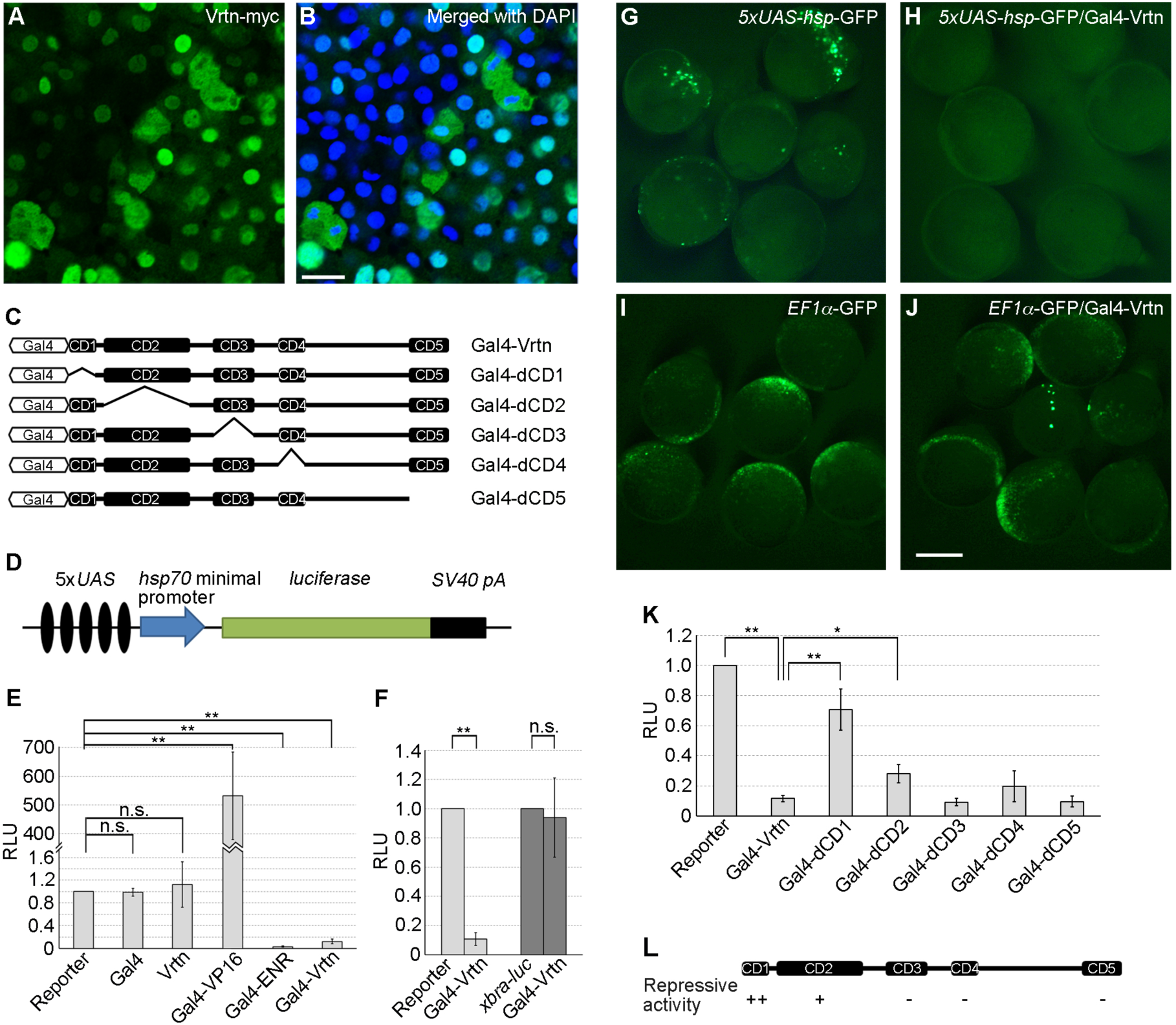
Vrtn functions as a transcriptional repressor. (A,B) Subcellular localization of Vrtn-myc in deep cells at 50% epiboly. (C) Full-length and truncated Vrtn proteins fused with Gal4 DNA-binding domain. (D) Schema of the *5xUAS-hsp-luc* reporter. (E) Graph shows the specificity and validity of the one-hybrid assay system. Bars represent the mean values ± s.d. from three independent experiments. (F) Gal4-Vrtn does not change luciferase activity driven by *xbra* promoter. Bars represent the mean values ± s.d. from three independent experiments. (G-J) Gal4-Vrtn suppresses GFP transcription driven by *5xUAS-hsp* promoter, but not by *EF1a* promoter, in WT embryos at bud stage. The embryos were injected with *5xUAS-hsp-GFP* plasmid (100 pg/embryo), *EF1a-GFP* plasmid (20 pg/embryo) alone or together with *Gal4-Vrtn* mRNA (200 pg/embryo). (K) Graph showing the transcriptional repressor activity of full-length and truncated Vrtn proteins. Bars represent the mean values ± s.d. from three independent experiments. (F) Summary of the repressive activity of different conserved domains. Scale bars: (A,B) 20 μm; (G-J) 200 μm.

We then used the UAS-Gal4 one-hybrid system to investigate how Vrtn functions as a transcriptional regulator (Fig. 6C), and synthetic mRNA (100 pg) was coinjected with a luciferase reporter driven by 5x UAS and a minimal *hsp70* promoter in zebrafish embryos (Fig. 6D). Gal4-Vrtn suppressed the reporter activity by almost 10 folds, in a similar manner as Gal4-ENR, a known transcriptional repressor. Gal4 or Vrtn alone had no effect on the reporter activity, and Gal4-Vrtn could not alter the activity of *xbra-luc*, an unrelated reporter (Fig. 6E,F). In addition, Gal4-Vrtn eliminated GFP expression under the control of UAS elements, but had no effect on GFP transcription driven by *EF1α* promoter (Fig. 6G-J). These demonstrate that Vrtn functions to repress gene transcription. We then tried to determine Vrtn functional domains involved in the repressor activity. Gal4-dCD1 only suppressed less than 30% of the luciferase activity, while Gal4-dCD2 showed a weak decrease of repressive activity (Fig. 6K,I). However, Gal4-dCD3, Gal4-dCD4 or Gal4-dCD5 retained a similar repressive activity as Gal4-Vrtn (Fig. 6K,L). These indicate that Vrtn acts as a potent transcriptional repressor and that its N-terminal domain is critical for the repressor function.

### Vrtn binds *bmp2b* upstream sequence and represses its transcription

We further examined whether Vrtn binds *bmp2b* regulatory region by chromatin immunoprecipitation followed by qPCR analysis (ChIP-qPCR). Zebrafish embryos were injected with synthetic *vrtn-myc* mRNA (100 pg) at 1-cell stage and cross-linked at 50% epiboly. The sonicated chromatins were precipitated with anti-myc antibody. Following qPCR analysis to detect DNA fragments of *bmp2b* gene (Fig. 7A), a -4775 to -4664 amplicon was significantly enriched by Vrtn-myc (Fig. 7B). To test if this region could act as a functional silencer of *bmp2b*, a -5701 to -3772 fragment and the promoter region (-800 to +605) were fused to luciferase reporter. A control reporter contained only the -800 to +605 promoter region (Fig. 7C). Synthetic *vrtn* mRNA (500 pg) was coinjected with these reporters at 1-cell stage and luciferase activity was assayed at 50% epiboly. We found that Vrtn suppressed luciferase activity by about three folds in the presence of the -5701 to -3772 fragment, but it had no effect on the control reporter (Fig. 7D). Taken together, our results suggest that Vrtn interacts with *bmp2b* regulatory sequence to repress its transcription.

**Figure 7.**
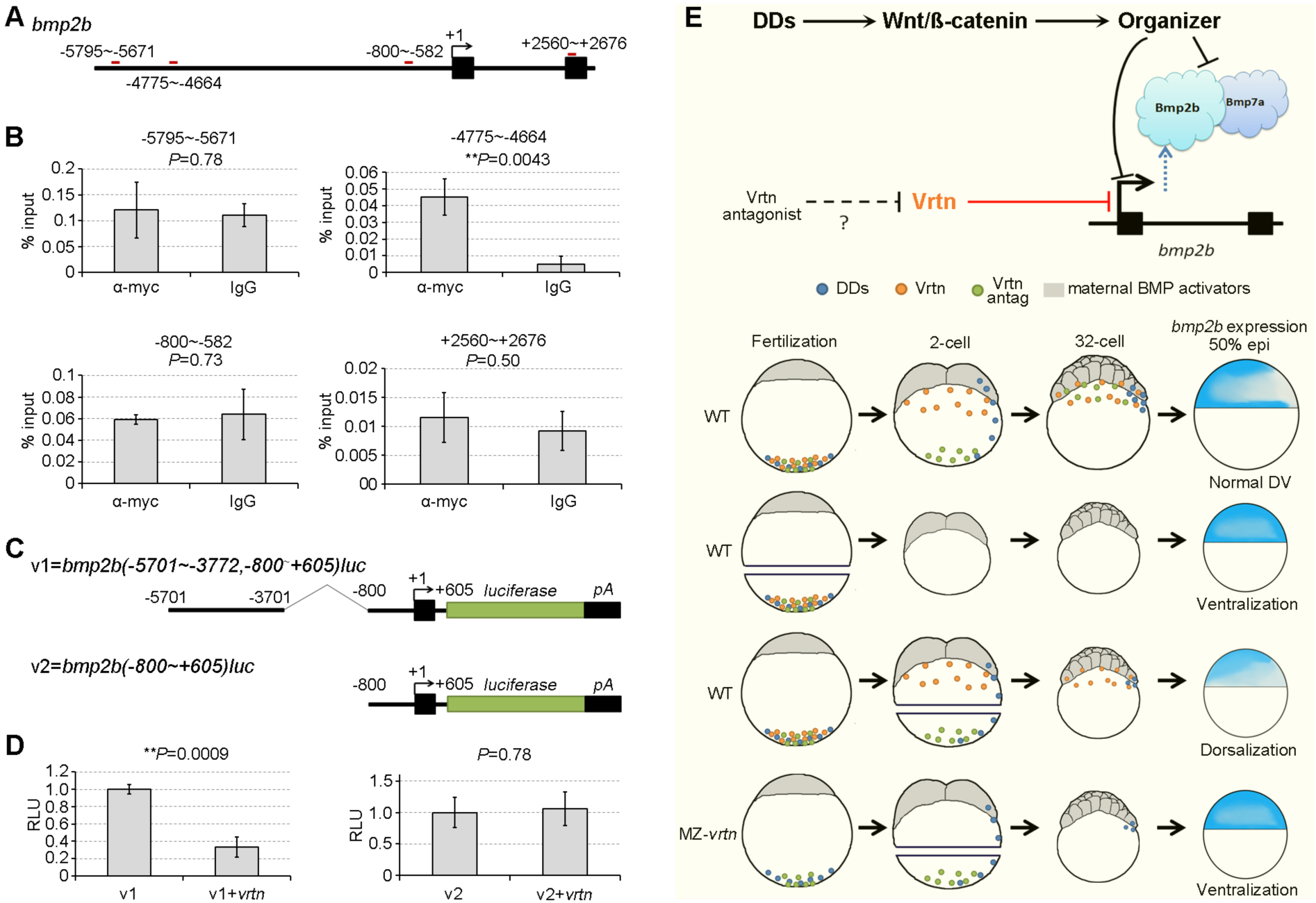
Vrtn binds to *bmp2b* upstream sequence and suppresses its transcription. (A) Location of amplicons (red lines) in *bmp2b* locus. (B) The -4775∼-4664 fragment is significantly enriched by anti-myc antibody. Bars represent the mean values ± s.d. from three independent experiments. (C) Representation of *bmp2b* luciferase reporters. (D) Vrtn represses the v1, but not v2, reporter activity. The value obtained with the reporter alone is set as 1, and bars represent the mean values ± s.d. from three independent experiments. (E) The “dual-factor” model. Vrtn and a putative antagonist are vegetally localized factors that functions independently of DDs and the Spemann organizer. Vrtn is translocated fast away towards the animal pole and functions to inhibit the *bmp2b* marginal transcription, while the antagonist is translocated relatively slowly and should activate BMP signalling. VAb@1cell enhances *bmp2b* expression and ventralization by ablating both Vrtn and DDs. VAb@2cell, however, results in deficient *bmp2b* transcription and dorsalization by removing the antagonist. In VAb@2cell *MZ-vrtn* embryos, the absence of Vrtn activity and partial loss of DDs enhance *bmp2b* expression and produce ventralization.

## DISCUSSION

We show that Vrtn is a novel vegetally localized regulator of *bmp2b* transcription in DV patterning during zebrafish early development. Maternal *vrtn* transcripts are actively transported to the animal pole region before the first cleavage and co-localizes with *bmp2b* in the margin after the onset of zygotic transcription. Vrtn interacts with *bmp2b* upstream regulatory region and represses its transcription independently of Wnt/ß-catenin signalling. These findings identify a novel maternal mechanism that ensures a correct BMP gradient for DV axis formation.

It is well established that the Spemann organizer expresses both BMP signalling antagonists and BMP ligands (Carron and Shi, 2016), which is required for establishing a correct BMP gradient (Xue et al., 2014), and represents a mechanism of self-regulation of the morphogenetic field (Reversade et al., 2005; De Robertis, 2006). Thus, both the expression, including the initial transcription and late maintenance, and the activity of BMP ligands in the Spemann organizer are tightly regulated by either a positive or a negative mechanism. There are several lines of evidence indicating that BMP activity in the ventral region is controlled by similar mechanisms. In zebrafish, *bmp2b, bmp7a* and *bmp4* are expressed in the lateral and ventral regions during late blastula and gastrula stages. Several maternal factors, including Gdf6a and Pou5f3, were shown to play a critical role for initiating their zygotic transcription (Sidi et al., 2003; Reim and Brand, 2006). However, it is unclear whether there are maternal factors preventing their excessive up-regulation before the formation of the Spemann organizer. In the present study, we uncovered Vrtn as a maternal factor representing one of the possible mechanisms limiting BMP activation. In particular, we find that Vrtn-induced dorsalization is achieved by transcriptional repression of *bmp2b* expression in the lateral and ventral margins of the late blastula.

The down-regulation of *bmp2b* by Vrtn is not a consequence on the positive auto-regulation of BMP signalling. In *swr* or *snh* mutants with loss-of-function of *bmp2b* or *bmp7a*, respectively, BMP signalling is blocked due to the absence of functional Bmp2b/Bmp7a heterodimer, which impairs *bmp2b* positive auto-regulation. However, in each mutant, *bmp2b* expression is only strongly reduced at mid- to late-gastrula stages, while it is almost normal at shield stage (Nguyen et al., 1998; Hild et al., 1999; Kondo, 2007). This indicates that *bmp2b* positive feedback loop is established after shield stage, much later than the initiation of its marginal up-regulation at 30% epiboly. Furthermore, *vrtn* overexpression only abolish *bmp2b* expression in the ventral ectoderm at mid-gastrula stage, leaving a relatively intact or enhanced expression in the I-YSL, while in *swr* or *snh* mutant, *bmp2b* expression shows an overall reduction, both in the ectoderm and I-YSL (Nguyen et al., 1998). These suggest that the marginal up-regulation of *bmp2b* transcription should not be controlled by the positive feedback loop of BMP signalling. It is likely activated by maternal factors such as Gdf6a and Pou5f3 (Sidi et al., 2003; Reim and Brand, 2006), while Vrtn plays an inhibitory role to limit this up-regulation.

The mechanism by which Vrtn functions in DV axis formation is also different from other reported maternal factors. The marginal transcription of *bmp2b* at shield stage is intact in *runx2bt2* morphant, *MZp18ahub* (*ints6*) and MZ*split top* (*ctsba*) mutant embryos (Flores et al., 2008; Kapp et al., 2013; Langdon et al., 2016). Loss-of-function of these genes, as well as *lnx2b* (Ro and Dawid, 2009), leads to an early expansion of Spemann organizer genes expression, often associated with a down-regulation of *ved* and *vox*, and/or reduced zygotic Wnt/ß-catenin signalling. The detailed molecular mechanisms by which these genes regulate DV axis formation are not known, except for Lnx2b, which is an E3 ubiquitin ligase that negatively regulates Bozozok function through ubiquitin proteasome pathway (Ro and Dawid, 2009). Nevertheless, the above phenotypes are obviously different from those resulted from VAb@2cell and Vrtn overexpression, which do not seem to cause a reduction of zygotic Wnt/ß-catenin activity and an early expansion of *gsc* and *chd*. These data strongly suggest that Vrtn functions in parallel with the presently known maternal DV regulators.

Both Vrtn overexpression and VAb@2cell similarly inhibit *bmp2b* marginal transcription at shield stage and restrict its expression only in I-YSL during gastrulation. This situation is highly similar to that in *MZsmad5* mutant (Hild et al., 1999; Kondo, 2007), and further supports that Vrtn functions specifically to regulate BMP signalling. However, unlike *MZsmad5* mutant, cells from VAb@2cell embryos are capable of interpreting BMP signal, suggesting that Vrtn is not a component of the BMP pathway, but it functions as a regulator for *bmp2b* transcription. Our data show that Vrtn binds to the -4775 to -4664 region upstream of *bmp2b* promoter where no Smad-binding motif can be predicted. Thus, it is likely that Vrtn represses Smad5-mediated *bmp2b* transcription at the margin. Further studies are needed to test this possibility.

It should be mentioned that Vrtn alone would not be enough to account for the VAb@2cell-induced dorsalization. In particular, how does the repressor function of Vrtn become activated in VAb@2cell embryos? Why *vrtn* mutant displays a mildly ventralized phenotype? Whether there are BMP activators removed after VAb@2cell? Although these questions are still open for further investigation, one likely explanation may be presence of another vegetally localized putative BMP activator, or Vrtn antagonist, which is involved in the activation or “de-repression” of BMP signalling in the margin and ventral ectoderm during gastrula stages. This factor may move relatively slowly after fertilization, and its transport would persist during early cleavage stages. Supporting this hypothesis, the dorsalizing effect of Vrtn can be antagonized by vegetal RNA at 2- to 4-cell stages, but the Vrtn-antagonizing activity of vegetal RNA declines at later stages. Taking these results into consideration, we propose a "dual-factor" model to explain Vrtn function in DV patterning. In normal situation, Vrtn and the BMP activator mutually antagonize, either directly or indirectly, to establish a BMP signalling gradient. In VAb@1cell embryos, all vegetal factors are removed, including DDs, Vrtn and the BMP activator. In the absence of inhibition, *bmp2b* expression is enhanced and the embryos are ventralized. By contrast, in VAb@2cell embryos, most of the BMP activator and part of DDs are removed, but sufficient amounts of Vrtn are transported to the blastoderm and freed from its antagonist. This leads to inhibition of BMP signalling and dorsalization, even though the activity of DDs may be reduced (Fig. 7E). Supporting this hypothesis, VAb@2cell is unable to dorsalize MZ*vrtn* embryos due to the absence of Vrtn. In this situation, the ablated MZ*vrtn* embryos only maintain a reduced amount of DDs, similarly as VAb@1cell WT embryos. Thus, it is not surprising that VAb@2cell MZ*vrtn* embryos exhibit an increased *bmp2b* expression and severely ventralized phenotype.

In conclusion, Vrtn is novel maternal transcriptional repressor that modulates *bmp2b* expression to maintain a correct BMP gradient along the DV axis. It acts independently of the Spemann organizer to prevent an excess of ventral fate in the early embryo. To further decipher the mechanism by which it regulates DV patterning, it will be interesting to identify potential Vrtn-associated events such as the presence and activity of Vrtn-antagonizing factors and interacting proteins.

## MATERIALS AND METHODS

### VAb

Ablation of vegetal yolk at different stages was performed essentially as described (Mizuno et al., 1999).

### Transplantation

The embryos were injected with RLDx, and labelled cells from the animal pole region at sphere stage were transplanted to the margin of unlabeled recipients in Ringer’s without egg albumen. Chimeric embryos were cultured in 1/3 x Ringer’s with 1.6% egg albumen and antibiotics, and fixed at appropriate stages in a Petri dish with 4% paraformaldehyde in Ringer’s overnight at room temperature. For transplantation of mixed cells, synchronously fertilized eggs were labelled with FLDx or RLDx. At sphere stage, 20 blastoderms from FLDx-labelled intact and RLDx-labelled embryos were treated with trypsin-EDTA solution (2 mg/ml trypsin, 1 mM EDTA in PBS) at 37°C for 15 minutes, and dissociated into single cells by gentle pipetting. FLDx- and RLDx-labelled cells were thoroughly mixed and collected by centrifugation at 250 g. The cell pellet was suspended in Ringer’s and a small fraction of mixed cells was microinjected into the ventral margin of unlabeled WT recipients at shield stage.

### Microinjections

The coding regions of *bmp2b, vrtn, dazl, celf1* and *magoh* were cloned in pCS2 vector, and capped mRNAs were in vitro transcribed using mMESSAGE mMACHINE SP6 kit (Ambion). Dechorionated embryos were injected in the blastoderm at 1-cell stage. The bmp2bMO (Lele et al., 2001) and *ß-catenin2MO* (Bellipanni et al., 2006) were synthesized by Gene Tools.

### Extraction of vegetal RNA

*MZvrtn* embryos were dechorionated and the animal halves were removed. The remaining vegetal explants were immediately transferred to Trizol reagent (Invitrogen) for RNA extraction. For each stage, total RNA from 50 vegetal explants was dissolved in 3 μl RNase-free water. An equal volume of synthetic *vrtn* mRNA (100 ng/μl) and the vegetal RNA was thoroughly mixed, and a mixture (5 nl) was injected into each embryo.

### RNA-seq

Total RNA was isolated from animal and vegetal pole regions dissected at 1-cell stage. The mRNA libraries were constructed by the TruSeq RNA Library Preparation Kit, and 100 bp paired end sequencing was performed on Illumina HiSeq 2000. RNA-Seq data was aligned to reference transcriptome assembly (www.ensembl.org), and analyzed using EdgeR software.

### CRISPR/Cas9-mediated genome editing

Capped mRNA for zebrafish codon-optimized Cas9 (Liu et al., 2014) was synthesized using mMESSAGE mMACHINE T7 kit (Ambion). The *vrtn* targeting sequence was cloned into p-T7-gRNA vector, and the DNA template for target-specific gRNA was PCR amplified as described (Xiao et al., 2013).

### ISH and qRT-PCR

Antisense RNA probes labelling and ISH was carried out as described (Shao et al., 2012). Total RNA was extracted using Trizol reagent and treated with with DNase I. They were reverse transcribed using M-MLV reverse transcriptase (Invitrogen), and qPCR with specific primers (Table S2) was performed using Quant qRT-PCR kit (Tiangen). The 2^-ΔΔCt^ method was employed to estimate relative expression level.

### Luciferase assays and ChIP-qPCR

Synthetic mRNA was coinjected with reporter DNA at 1-cell stage and 15 embryos at 70% epiboly were lysed and assayed using the Dual-Luciferase^®^ Reporter Assay System (Promega). ChIP-qPCR was carried out as described (Bogdanovic et al., 2013).

### Statistical analysis

The unpaired Student’s *t*-test was used and *P* values less than 0.05 and 0.01 were considered as significant (*) and extremely significant (**), respectively.

## Acknowledgements

We thank F. Liu, A.M. Meng for reagents, the members of our laboratory for zebrafish care, and the imaging facility at the School of life science (Shandong University) for confocal image acquisition.

## Author contributions

M.S. and D.-L.S. conceived and designed the experiments. M.S. performed VAb and transplantation experiments. M.W. and M.S. realised knockout of *vrtn*, and other molecular and biochemical analysis. M.S., Y.-W.G. and M.W. identified vrtn. M.S., Y.-J.Z., and D.-L.S. analyzed the data. M.S. and D.-L.S. wrote the paper.

## References

Abrams, E. W. and Mullins, M.C. (2009). Early zebrafish development: it's in the maternal genes. Curr. Opin. Genet. Dev. 19, 396–403.

Bellipanni, G., Varga, M., Maegawa, S., Imai, Y., Kelly, C., Myers, A. P., Chu, F., Talbot, W. S. and Weinberg, E. S. (2006). Essential and opposing roles of zebrafish beta-catenins in the formation of dorsal axial structures and neurectoderm. Development 133, 1299–1309.

Bogdanovic, O., Fernandez-Minan, A., Tena, J. J., de la Calle-Mustienes, E. and Luis Gomez-Skarmeta, J. (2013). The developmental epigenomics toolbox: ChIP-seq and MethylCap-seq profiling of early zebrafish embryos. Methods 62, 207–215.

Carron, C. and Shi, D. L. (2016). Specification of anteroposterior axis by combinatorial signaling during Xenopus development. Wiley Interdiscip. Rev. Dev. Biol. 5, 150–168.

Cuykendall, T. N. and Houston, D. W. (2009). Vegetally localized Xenopus trim36 regulates cortical rotation and dorsal axis formation. Development 136, 3057–6305.

De Robertis, E. M. (2006). Spemann's organizer and self-regulation in amphibian embryos. Nat. Rev. Mol. Cell. Biol. 7, 296–302.

Dick, A., Hild, M., Bauer, H., Imai, Y., Maifeld, H., Schier, A. F., Talbot, W. S., Bouwmeester, T. and Hammerschmidt, M. (2000). Essential role of Bmp7 (snailhouse) and its prodomain in dorsoventral patterning of the zebrafish embryo. Development 127, 343–354.

Dosch, R. and Niehrs, C. (2000). Requirement for anti-dorsalizing morphogenetic protein in organizer patterning. Mech. Dev. 90, 195–203.

Flores, M. V., Lam, E. Y., Crosier, K. E. and Crosier, P. S. (2008). Osteogenic transcription factor Runx2 is a maternal determinant of dorsoventral patterning in zebrafish. Nat. Cell. Biol. 10, 346–352.

Ge, X., Grotjahn, D., Welch, E., Lyman-Gingerich, J., Holguin, C., Dimitrova, E., Abrams, E. W., Gupta, T., Marlow, F. L., Yabe, T., et al. (2014). Hecate/Grip2a acts to reorganize the cytoskeleton in the symmetry-breaking event of embryonic axis induction. PLoS Genet. 10, e1004422.

Goutel, C., Kishimoto, Y., Schulte-Merker, S. and Rosa, F. (2000). The ventralizing activity of Radar, a maternally expressed bone morphogenetic protein, reveals complex bone morphogenetic protein interactions controlling dorso-ventral patterning in zebrafish. Mech. Dev. 99, 15–27.

Hild, M., Dick, A., Rauch, G. J., Meier, A., Bouwmeester, T., Haffter, P. and Hammerschmidt, M. (1999). The smad5 mutation somitabun blocks Bmp2b signaling during early dorsoventral patterning of the zebrafish embryo. Development 126, 2149–2159.

Inui, M., Montagner, M., Ben-Zvi, D., Martello, G., Soligo, S., Manfrin, A., Aragona, M., Enzo, E., Zacchigna, L., Zanconato, F., et al. (2012). Self-regulation of the head-inducing properties of the Spemann organizer. Proc. Natl. Acad. Sci. U. S. A. 109, 15354–15359.

Jesuthasan, S. and Stahle, U. (1997). Dynamic microtubules and specification of the zebrafish embryonic axis. Curr. Biol. 7, 31–42.

Kapp, L. D., Abrams, E. W., Marlow, F. L. and Mullins, M. C. (2013). The integrator complex subunit 6 (Ints6) confines the dorsal organizer in vertebrate embryogenesis. PLoS Genet. 9, e1003822.

Kishimoto, Y., Lee, K. H., Zon, L., Hammerschmidt, M. and Schulte-Merker, S. (1997). The molecular nature of zebrafish swirl: BMP2 function is essential during early dorsoventral patterning. Development 124, 4457–4466.

Kondo, M. (2007). Bone morphogenetic proteins in the early development of zebrafish. FEBS J. 274, 2960–2967.

Langdon, Y. G., Fuentes, R., Zhang, H., Abrams, E. W., Marlow, F. L. and Mullins, M.C. (2016). Split top: a maternal cathepsin B that regulates dorsoventral patterning and morphogenesis. Development 143, 1016–1028.

Langdon, Y. G. and Mullins, M. C. (2011). Maternal and Zygotic Control of Zebrafish Dorsoventral Axial Patterning. Annu. Rev. Genet. 45, 357–377.

Lele, Z., Bakkers, J. and Hammerschmidt, M. (2001). Morpholino phenocopies of the swirl, snailhouse, somitabun, minifin, silberblick, and pipetail mutations. Genesis 30, 190–194.

Leung, T., Bischof, J., Soll, I., Niessing, D., Zhang, D., Ma, J., Jackle, H. and Driever, W. (2003). bozozok directly represses bmp2b transcription and mediates the earliest dorsoventral asymmetry of bmp2b expression in zebrafish. Development 130, 3639–3649.

Little, S. C. and Mullins, M. C. (2009). Bone morphogenetic protein heterodimers assemble heteromeric type I receptor complexes to pattern the dorsoventral axis. Nat. Cell Biol. 11, 637–643.

Liu, D., Wang, Z., Xiao, A., Zhang, Y., Li, W., Zu, Y., Yao, S., Lin, S. and Zhang, B. (2014). Efficient gene targeting in zebrafish mediated by a zebrafish-codon-optimized cas9 and evaluation of off-targeting effect. J. Genet. Genomics 41, 43–46.

Lu, F. I., Thisse, C. and Thisse, B. (2011). Identification and mechanism of regulation of the zebrafish dorsal determinant. Proc. Natl. Acad. Sci. U. S. A. 108, 15876–15880.

Maegawa, S., Yasuda, K. and Inoue, K. (1999). Maternal mRNA localization of zebrafish DAZ-like gene. Mech. Dev. 81, 223–226.

Marlow, F. L. and Mullins, M. C. (2008). Bucky ball functions in Balbiani body assembly and animal-vegetal polarity in the oocyte and follicle cell layer in zebrafish. Dev. Biol. 321, 40–50.

Mei, W., Lee, K. W., Marlow, F. L., Miller, A. L. and Mullins, M. C. (2009). hnRNP I is required to generate the Ca2+ signal that causes egg activation in zebrafish. Development 136, 3007–3017.

Mikawa, S., Sato, S., Nii, M., Morozumi, T., Yoshioka, G., Imaeda, N., Yamaguchi, T., Hayashi, T. and Awata, T. (2011). Identification of a second gene associated with variation in vertebral number in domestic pigs. BMC Genet. 12, 5.

Miller, J. R., Rowning, B. A., Larabell, C. A., Yang-Snyder, J. A., Bates, R. L. and Moon, R. T. (1999). Establishment of the dorsal-ventral axis in Xenopus embryos coincides with the dorsal enrichment of dishevelled that is dependent on cortical rotation. J. Cell Biol. 146, 427–437.

Mizuno, T., Yamaha, E., Kuroiwa, A. and Takeda, H. (1999). Removal of vegetal yolk causes dorsal deficencies and impairs dorsal-inducing ability of the yolk cell in zebrafish. Mech. Dev. 81, 51–63.

Nguyen, V. H., Schmid, B., Trout, J., Connors, S. A., Ekker, M. and Mullins, M. C. (1998). Ventral and lateral regions of the zebrafish gastrula, including the neural crest progenitors, are established by a bmp2b/swirl pathway of genes. Dev. Biol. 199, 93–110.

Nojima, H., Rothhamel, S., Shimizu, T., Kim, C. H., Yonemura, S., Marlow, F. L. and Hibi, M. (2010). Syntabulin, a motor protein linker, controls dorsal determination. Development 137, 923–933.

Ober, E. A. and Schulte-Merker, S. (1999). Signals from the yolk cell induce mesoderm, neuroectoderm, the trunk organizer, and the notochord in zebrafish. Dev. Biol. 215, 167–181.

Ramel, M. C., Buckles, G. R., Baker, K. D. and Lekven, A. C. (2005). WNT8 and BMP2B co-regulate non-axial mesoderm patterning during zebrafish gastrulation. Dev. Biol. 287, 237–248.

Reim, G. and Brand, M. (2006). Maternal control of vertebrate dorsoventral axis formation and epiboly by the POU domain protein Spg/Pou2/Oct4. Development 133, 2757–2770.

Reversade, B., Kuroda, H., Lee, H., Mays, A. and De Robertis, E. M. (2005). Depletion of Bmp2, Bmp4, Bmp7 and Spemann organizer signals induces massive brain formation in Xenopus embryos. Development 132, 3381–3392.

Ro, H. and Dawid, I. B. (2009). Organizer restriction through modulation of Bozozok stability by the E3 ubiquitin ligase Lnx-like. Nat. Cell. Biol. 11, 1121–1127.

Shao, M., Lin, Y., Liu, Z., Zhang, Y., Wang, L., Liu, C. and Zhang, H. (2012). GSK-3 activity is critical for the orientation of the cortical microtubules and the dorsoventral axis determination in zebrafish embryos. PLoS One 7, e36655.

Sidi, S., Goutel, C., Peyriéras, N. and Rosa, F. M. (2003). Maternal induction of ventral fate by zebrafish radar. Proc. Natl. Acad. Sci. U. S. A. 100, 3315–3320.

Slack, J. (2014). Establishment of spatial pattern. Wiley Interdiscip. Rev. Dev. Biol. 3, 379–88.

Stickney, H. L., Imai, Y., Draper, B., Moens, C. and Talbot, W. S. (2007). Zebrafish bmp4 functions during late gastrulation to specify ventroposterior cell fates. Dev. Biol. 310, 71–84.

Sun, Y., Tseng, W. C., Fan, X., Ball, R. and Dougan, S. T. (2014). Extraembryonic signals under the control of MGA, Max, and Smad4 are required for dorsoventral patterning. Dev. Cell 28, 322–334.

Suzuki, H., Maegawa, S., Nishibu, T., Sugiyama, T., Yasuda, K. and Inoue, K. (2000). Vegetal localization of the maternal mRNA encoding an EDEN-BP/Bruno-like protein in zebrafish. Mech. Dev. 93, 205–209.

Tao, Q., Yokota, C., Puck, H., Kofron, M., Birsoy, B., Yan, D., Asashima, M., Wylie, C. C., Lin, X. and Heasman, J. (2005). Maternal wnt11 activates the canonical wnt signaling pathway required for axis formation in Xenopus embryos. Cell 120, 857–871.

Tran, L. D., Hino, H., Quach, H., Lim, S., Shindo, A., Mimori-Kiyosue, Y., Mione, M., Ueno, N., Winkler, C., Hibi, M., et al. (2012). Dynamic microtubules at the vegetal cortex predict the embryonic axis in zebrafish. Development 139, 3644–3652.

Weaver, C., Farr, G. H., Pan, W., Rowning, B. A., Wang, J., Mao, J., Wu, D., Li, L., Larabell, C. A. and Kimelman, D. (2003). GBP binds kinesin light chain and translocates during cortical rotation in Xenopus eggs. Development 130, 5425–5436.

Xiao, A., Wang, Z., Hu, Y., Wu, Y., Luo, Z., Yang, Z., Zu, Y., Li, W., Huang, P., Tong, X., et al. (2013). Chromosomal deletions and inversions mediated by TALENs and CRISPR/Cas in zebrafish. Nucleic Acids Res. 41, e141.

Xue, Y., Zheng, X., Huang, L., Xu, P., Ma, Y., Min, Z., Tao, Q., Tao, Y. and Meng, A. (2014). Organizer-derived Bmp2 is required for the formation of a correct Bmp activity gradient during embryonic development. Nat. Commun. 5, 3766.

